# Trade-offs between antiseptic cytotoxicity and efficacy in a human *ex vivo* wound contamination model

**DOI:** 10.1101/2021.02.14.430155

**Authors:** J. Z. Alex Cheong, Aiping Liu, Clayton J. Rust, Lindsay R. Kalan, Angela L. F. Gibson

## Abstract

Wound cleansing agents are routine in wound care, even in the absence of signs of infection. Antiseptic activity prevents contaminating microbes from establishing an infection while also raising concerns of cytotoxicity and delayed wound healing. Here, we used an *ex vivo* human skin excisional wound model to evaluate the cytotoxicity of five clinically-used wound cleaning agents (saline, povidone iodine, Dove® soap, Dial® soap, and chlorhexidine gluconate). We established a wound contamination model using ∼100 cells of *Pseudomonas aeruginosa* per wound to evaluate antiseptic efficacy and microbial biofilm spatial organization. We found that Dial® soap and chlorhexidine gluconate significantly reduced metabolic activity of the biopsies, while all treatments except saline affected local cellular viability. Within the contamination model, only chlorhexidine gluconate treatment resulted in significantly lower *P. aeruginosa* counts at 24 hours post-treatment, driven by sub-limit-of-detection counts immediately post-treatment. Later applications of chlorhexidine gluconate had no effects on microbial growth, with microscopy showing extensive surface colonization of the wound bed. We present a clinically-relevant model for evaluating antiseptic cytotoxicity and efficacy, with the ability to resolve spatial localization and temporal dynamics of tissue viability and microbial growth.

## INTRODUCTION

Microbial colonization and development of biofilm is hypothesized to contribute to impaired wound healing and excess inflammation (Azevedo et al., 2020; Rahim et al., 2017; Wolcott, 2017). To prevent microbial colonization and subsequent infection of wound tissue, wound cleansing agents ranging from saline to solutions of antiseptic compounds are routinely employed (Cambiaso-Daniel et al., 2018; Percival et al., 2016; Slaviero et al., 2018). Antimicrobial efficacy is a major consideration in the type of cleansing agent chosen, however antiseptics with high antimicrobial activity can be cytotoxic and may further inhibit tissue repair (Atiyeh et al., 2009; Barreto et al., 2020; Punjataewakupt et al., 2019; Smith, 2005; Thomas et al., 2009).

Classical models of evaluating antiseptic efficacy and cytotoxicity utilize reductionist models that evaluate components of human skin and wounds individually, such as through pure cell cultures or microbial cultures *in vitro* (Blenkharn, 1987; Borenfreund and Puerner, 1985; Cooper et al., 1991, 1990; Damour et al., 1992; Karpi and Sciences, 2017; Kloth et al., 2007; McLaughlin et al., 2020; Muller and Kramer, 2008; Niedner, 1997; Ponec et al., 1990; Rabenberg et al., 2002; Rembe et al., 2016; Tatuall et al., 1990). Importantly, such models lack the structural and compositional heterogeneity of human skin and spatiotemporal nature of wound tissue. *In vitro* models are unsuitable to evaluate clinical scenarios mimicking microbial contamination and progression of infection concurrently while evaluating antiseptic efficacy and cytotoxicity (Roberts et al., 2015). There is a critical need for clinically-applicable models that can simultaneously evaluate antimicrobial efficacy and cytotoxicity of antimicrobial interventions under a clinically-relevant context (Besser et al., 2020; Roche et al., 2019).

Here, we use an *ex vivo* human skin excisional wound model to evaluate the cytotoxicity of five clinically-used wound cleaning agents (saline, povidone iodine [PVI], Dove® soap, Dial® soap, and chlorhexidine gluconate [CHG]). To evaluate the antimicrobial efficacy of each wound cleanser, we established a wound contamination model by colonizing excisional wounds with ∼100 cells of *Pseudomonas aeruginosa* per wound and evaluated the expansion and biofilm formation of *P. aeruginosa* during topical treatment with each cleanser. To highlight the ability of our model to evaluate spatial and temporal dynamics, we use histological staining of cellular viability via lactate dehydrogenase (LDH) activity and microscopy to localize cytotoxicity and microbial colonization of the wound bed. We find that antiseptic cytotoxicity is correlated with antiseptic efficacy at early timepoints. Further, we show that CHG efficacy against *P. aeruginosa* is time-sensitive. Within 24 h of wound contamination, we find a significant 3-log reduction in *P. aeruginosa* total viable cell counts. However, with subsequent application of CHG after 24 h, diminished antiseptic activity is observed.

## RESULTS

### *Establishment of an ex vivo* wound contamination model

*Ex vivo* human skin wounds consisting of 6 mm partial thickness wounds on 12 mm full thickness skin biopsies were cultured in individual wells of a 12-well tissue culture plate in 3 mL of DMEM + 10% fetal bovine serum (FBS) with 0.15% agarose to support the biopsy at the air-media interface (Fig. 1a). We developed a contamination model to test the effects of clinically-used wound cleansing solutions under conditions where an infection was not suspect. Each wound was inoculated with ∼100 colony forming units (CFU) of *P. aeruginosa*. After 24 h of growth, 8.4 ± 0.5 (mean ± SD) log_10_ CFU/bisect of bacteria were recovered, a 6-log increase (Fig. 1b). Viable bacteria cells were not recovered above the limit of detection (50 CFU) in uninoculated wounds throughout the duration of the study (Fig. 1b).

**Figure 1.**
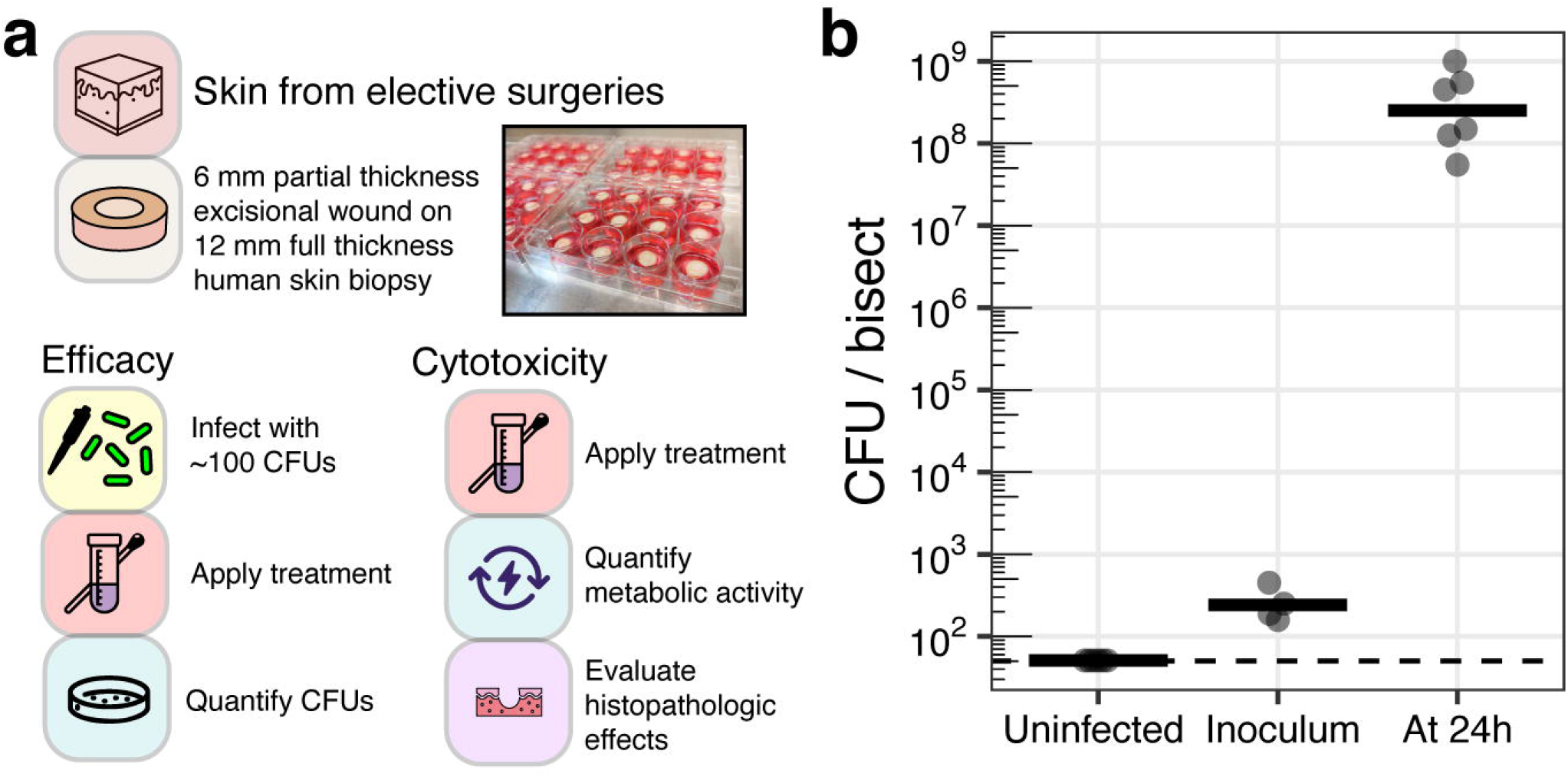
*Ex vivo* human excisional wound model to investigate antiseptic efficacy and cytotoxicity. (a) Overview of model. (b)Establishment of infection with ∼100 cells of *Pseudomonas aeruginosa* per wound to represent wound contamination. Biopsies were incubated for 24 h before enumeration. Uninfected wounds remained sterile with no microbial growth. Each data point represents one replicate bisect of a biopsy for uninfected and 24 h samples; horizontal bars show means of ≥ 6 replicates from ≥ 3 skin donors. For the inoculum, each data point represents a single quantification of an inoculum from ≥ 4 biological replicates.

### CHG effectively reduces bacteria bioburden at 24 hpt

The effectiveness of five clinically-used wound cleansing agents was tested. The cleansers including phosphate-buffered saline (PBS), povidone iodine (PVI), Dove® soap, Dial® soap, and chlorhexidine gluconate (CHG). Excisional wounds were inoculated in the contamination model and each antiseptic was applied topically for 30 min at 4 hours post-inoculation (hpi). At the time of treatment, no bacterial growth was observed, mimicking a clinical scenario warranting normal daily wound cleansing. Conversely, signs of infection such as local inflammation, visible slough, and bacterial growth would warrant an escalation of treatment. Quantitative viable cell counts at 24 h post-treatment (hpt) showed that CHG resulted in a significant ∼3 log_10_ CFU reduction of bacterial bioburden (*p. adj* < 0.0001, Fig. 2a) compared to an untreated control and the four other treatments.

**Figure 2.**
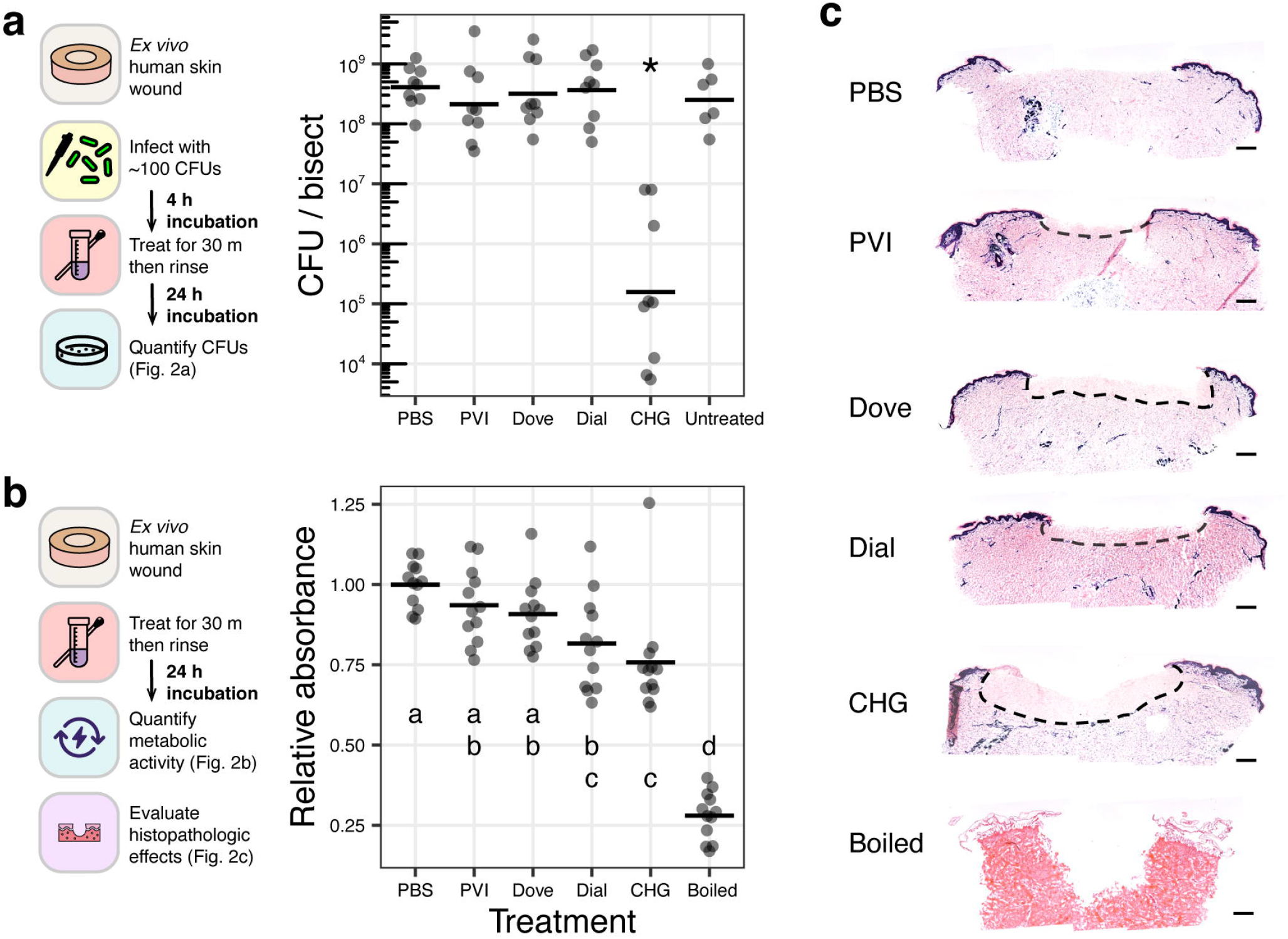
Antiseptic efficacy and cytotoxicity of topical wound cleansers. *Ex vivo* human excisional wound biopsies were evaluated at 24 h post-treatment. (a) Quantification of viable bacterial cell counts. Each data point represents one replicate bisect of a biopsy; horizontal bars show means of ≥ 9 replicates (for treatments) and ≥ 6 replicates (for untreated controls) from ≥ 3 skin donors. *, *p adj*. < 0.0001. (b) Quantification of metabolic activity relative to mean of PBS treatment using the MTT assay. Boiled treatment is negative viability control. Each data point represents one replicate biopsy; horizontal bars show means of ≥ 12 replicates from ≥ 4 skin donors. Treatments marked with different letters are significantly different from one another; *p adj*. < 0.05. (c) Histopathological assessment of cellular viability using LDH staining of 5 µm cryosections. Dark blue stain indicates viable cells. Dashed lines demarcate regions of depleted cellular viability. Scale bars represent 500 μm. Micrographs are representative of ≥ 3 skin donors.

### Wound cleansing agents have varying levels of cytotoxicity

To investigate the cytotoxicity of the antiseptics, the *ex vivo* excisional wound biopsies were treated with PBS, PVI, Dove® soap, Dial® soap, and CHG for 30 min in the absence of *P. aeruginosa*. At 24 hpt, we used the MTT assay to measure tissue metabolism as a surrogate for cell viability. We found that CHG and Dial® soap treatments resulted in a significant reduction of tissue metabolism compared to PBS-treated biopsies (*p. adj* < 0.05; Fig. 2b). In particular, the metabolic level in the CHG-treated tissue biopsies was also significantly lower than that of PVI-and Dove® soap-treated biopsies (*p. adj* < 0.05; Fig. 2b). Tissue viability was evaluated histologically by staining for lactate dehydrogenase (LDH) activity. A region of depleted cellular viability was identified in the epidermis at the wound edges and in the mid-reticular dermis of the wound in CHG-treated tissue biopsies, whereas loss of cell viability was localized more superficially in the dermis of the wound in Dial® and Dove® soap-treated biopsies (Fig. 2c). Both the MTT assay and LDH staining indicate that CHG is more cytotoxic on human skin than the other antiseptics (Fig. 2b, c).

### Temporal dynamics of CHG cytotoxicity and efficacy

Since our data showed that CHG has the highest antimicrobial efficacy and the greatest cytotoxicity within a 24 h timeframe, we wanted to determine if the cytotoxic effects persist over time. To evaluate this, we treated *ex vivo* wounds with either CHG or PBS for 30 min, followed by rinsing in sterile PBS, and then placed the tissue biopsies in culture for 14 days. At various time points, biopsies were harvested to assess cell viability. PBS-treated tissue biopsies remained viable up to 14 days, showing a dark blue stain indicative of LDH activity and cellular viability across the epidermis and throughout the dermis (Fig. 3c-f). Conversely, CHG treatment resulted in a progression of cytotoxicity (Fig. 3g-j). Consistent with early results, we observed at 1-day post-treatment loss of cellular viability at the epidermal wound edge and dermis where CHG was in direct contact with the tissue (Fig. 3g). On day 3, this loss of cellular viability progressed laterally from the wound across the epidermis (Fig. 3h). By day 7, cellular viability was lost from the entire 12mm biopsy (Fig. 3i, j), suggesting that despite rinsing away CHG after the 30 min treatment, cytotoxicity persists and leads to a profound progression of cellular injury.

**Figure 3.**
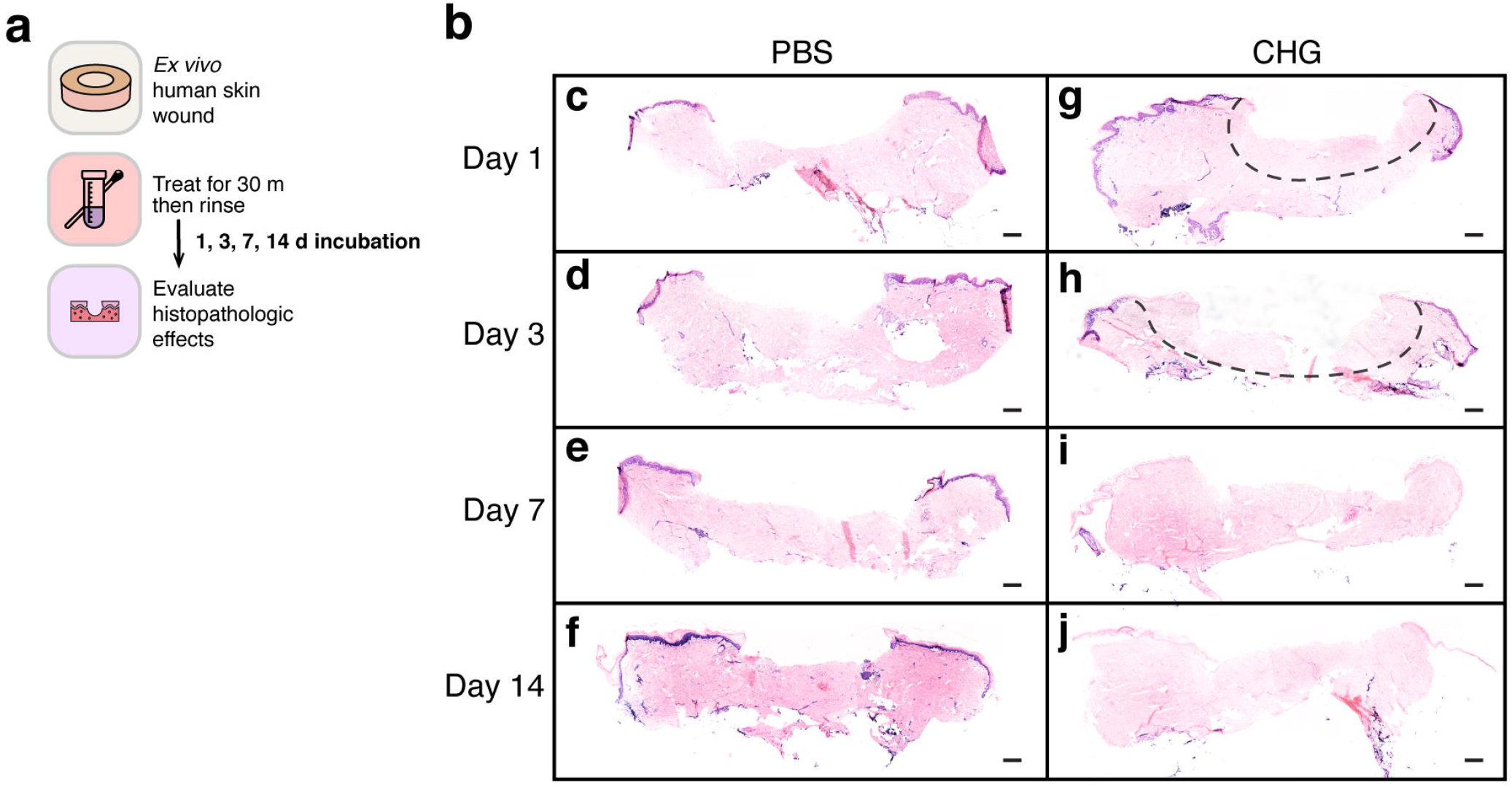
Temporal dynamics of CHG cytotoxicity. (a) Timeline of experiments. (b) Histopathological assessment of cellular viability using LDH staining of 5 um cryosections. Dark blue stain indicates viable cells. Dashed lines demarcate regions of depleted cellular viability. Scale bars represent 500 μm. Micrographs are representative of ≥ 3 skin donors. (c, d, e, f) Biopsies treated with PBS and processed at 1, 3, 7, and 14 days post-treatment respectively. (g, h, i, j) Biopsies treated with CHG and processed at 1, 3, 7, and 14 days post-treatment respectively.

Conversely, the antiseptic efficacy of CHG does not appear to be long lasting. We found CHG effectively reduces viable counts to below the limit-of-detection (50 CFUs) immediately post-application (≤ 1.7 vs. 2.4 ± 0.3 log_10_ CFU/bisect in PBS-treated wounds; *p* < 0.05; Fig. 4b), with effects lasting up to 24 hpt (5.2 ± 1.2 vs. 8.6 ± 0.3 log_10_ CFU/bisect in PBS-treated wounds; Fig. 2a). However, by 48 hpt, viable counts increased and were consistent with PBS-treated wounds (8.8 ± 0.2 vs. 9.0 ± 0.2 log_10_ CFU/bisect respectively; Fig. 4c). To mimic a once-daily clinical wound cleansing schedule, a second treatment of CHG was applied for 30 min to each contaminated wound 24 h after the first treatment, rinsed off, and incubated for another 24 h before processing. After the second CHG treatment, we observed viable counts that were not significantly different from a single CHG treatment at the same time point (8.5 ± 0.3 vs. 8.8 ± 0.2 log_10_ CFU/bisect in singly-treated wounds; Fig. 4c). CHG treatment applied at 24 h after cleansing with PBS also did not result in significantly different viable counts (8.9 ± 0.2 vs. 9.0 ± 0.2 log_10_ CFU/bisect in wounds singly-cleansed with PBS at 4 hpi; Fig. 4c).

**Figure 4.**
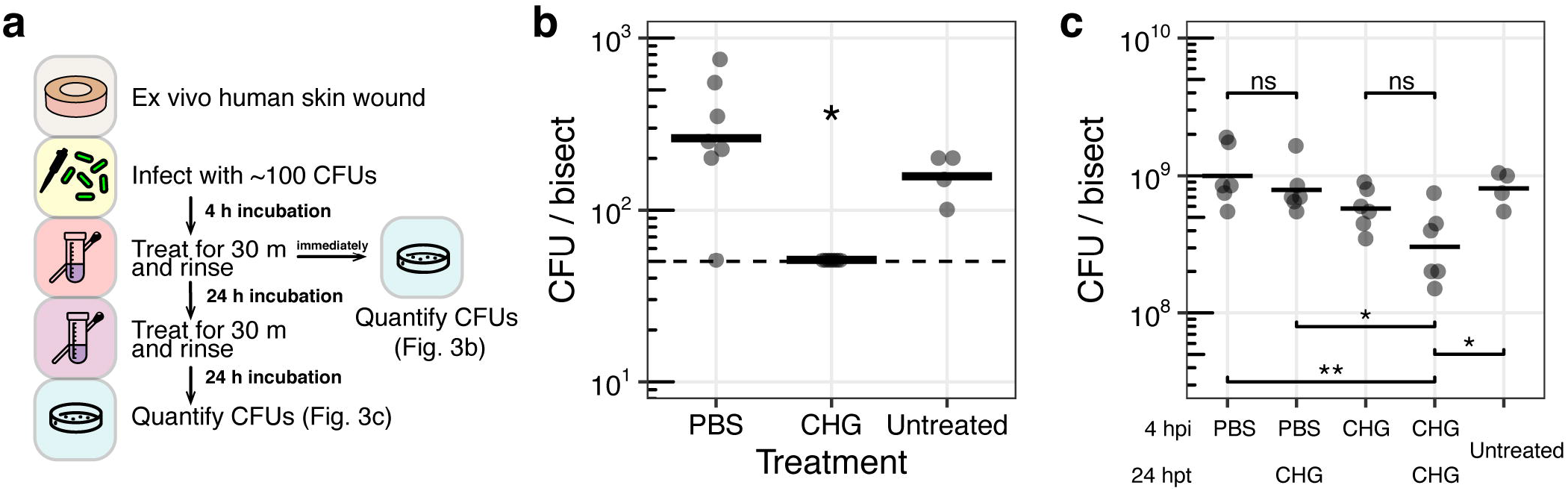
Temporal dynamics of CHG antiseptic efficacy. (a) Timeline of experiments. (b) *Ex vivo* human excisional wound biopsies were evaluated immediately post-treatment at 4 h post-infection. Dashed line indicates limit of detection (50 CFU). *, *p adj*. < 0.05. (c) *Ex vivo* human excisional wound biopsies were evaluated 24 h after second treatment. Treatments were applied at 4 h post-infection onto all wounds, and for a subset of wounds, again at 24 h post-treatment. Each data point represents one replicate bisect of a biopsy; horizontal bars show means of ≥ 6 replicates (for treatments) and ≥ 4 replicates (for untreated controls) from ≥ 2 skin donors. *, *p adj*. < 0.05; **, *p adj*. < 0.01; ns, not significantly different.

We then used scanning electron microscopy (SEM) to qualitatively evaluate biofilm architecture and surface topography of the wounds. In uninfected control wounds, the wound bed is a topologically heterogenous substrate for microbial attachment and growth (Fig. 5a). Large bundles of collagen fibers make up the connective tissue of the dermis, and are comprised of individual collagen fibrils, as shown by the clear banding pattern in the fibrils (Fig. 5a inset; Gottardi et al. 2016; Ushiki 2002). We found that colonized wounds treated with PBS at 4 hpi were covered with a dense layer of bacteria and extracellular matrix that completely obscures the collagen fibers of the wound bed (Fig. 5b). Bacterial cells were not detected on the surface of colonized wounds treated with CHG at 4 hpi (Fig. 5c), supporting our findings that CHG is efficacious up to 24 hpt. However, wounds treated with a second application of CHG at 24 hpt are again covered with a dense layer of bacterial cells and extracellular matrix over the wound bed (Fig. 5d), suggesting that CHG loses efficacy at later time points, in line with our culture data.

**Figure 5.**
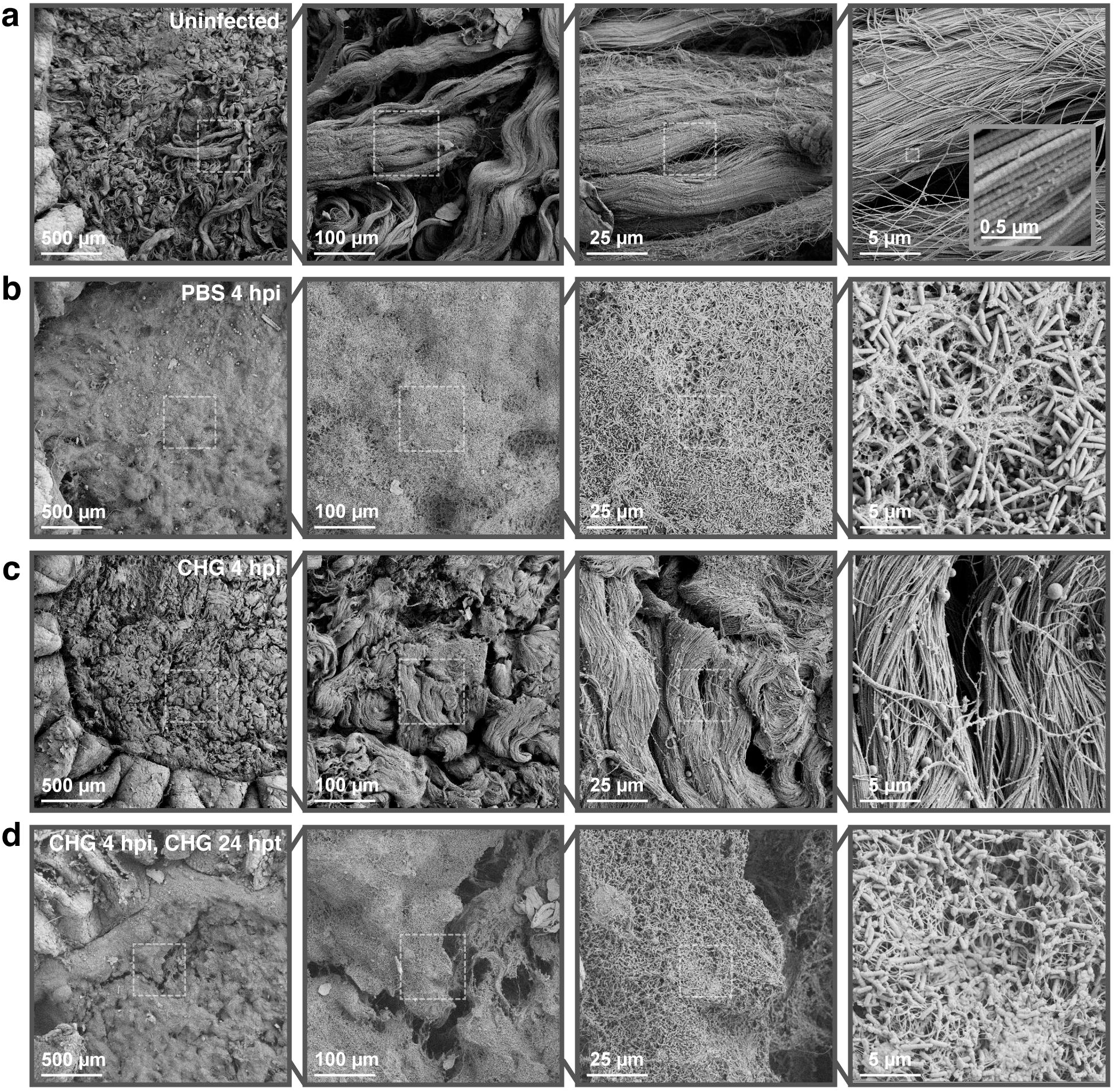
Scanning electron microscopy allows evaluation of microbial wound colonization. Scanning electron micrographs at four different magnifications (100x, 500x, 2 000x, 10 000x) of *ex vivo* wounds collected at 24 h after final treatment. Dashed outlines represent region magnified. (a) Uninfected wound. Inset micrograph shows banding pattern of collagen fibrils at 100 000x magnification. (b) PBS treatment at 4 hpi. (c) CHG treatment at 4 hpi (c) CHG treatment at 4 hpi, CHG treatment at 24 hpt.

We were intrigued by the lack of visible bacteria cells on wounds treated with CHG at 4 hpi (Fig. 5c), as these wounds had a bioburden of 5.2 ± 1.2 log_10_ CFU/bisect (Fig. 2a). As SEM shows only surface topology, we hypothesized that bacteria may be localized deeper in the tissue after treatment. We used confocal laser scanning microscopy (CLSM) for a depth-resolved perspective into the tissue. This technique revealed that wounds treated with CHG at 4 hpi contained single bacterial cells and small clusters deep within the wound bed and tissue (Fig. 6a), suggesting that although bacterial cells were not detected on the surface with SEM, migration into deeper tissues results in a reservoir within the wound to repopulate the wound surface. CLSM of wounds treated with PBS at 4 hpi and 24 hpi revealed large aggregates of *P. aeruginosa* (Fig. 6b), consistent with published biofilm models of this bacterial pathogen.

**Figure 6.**
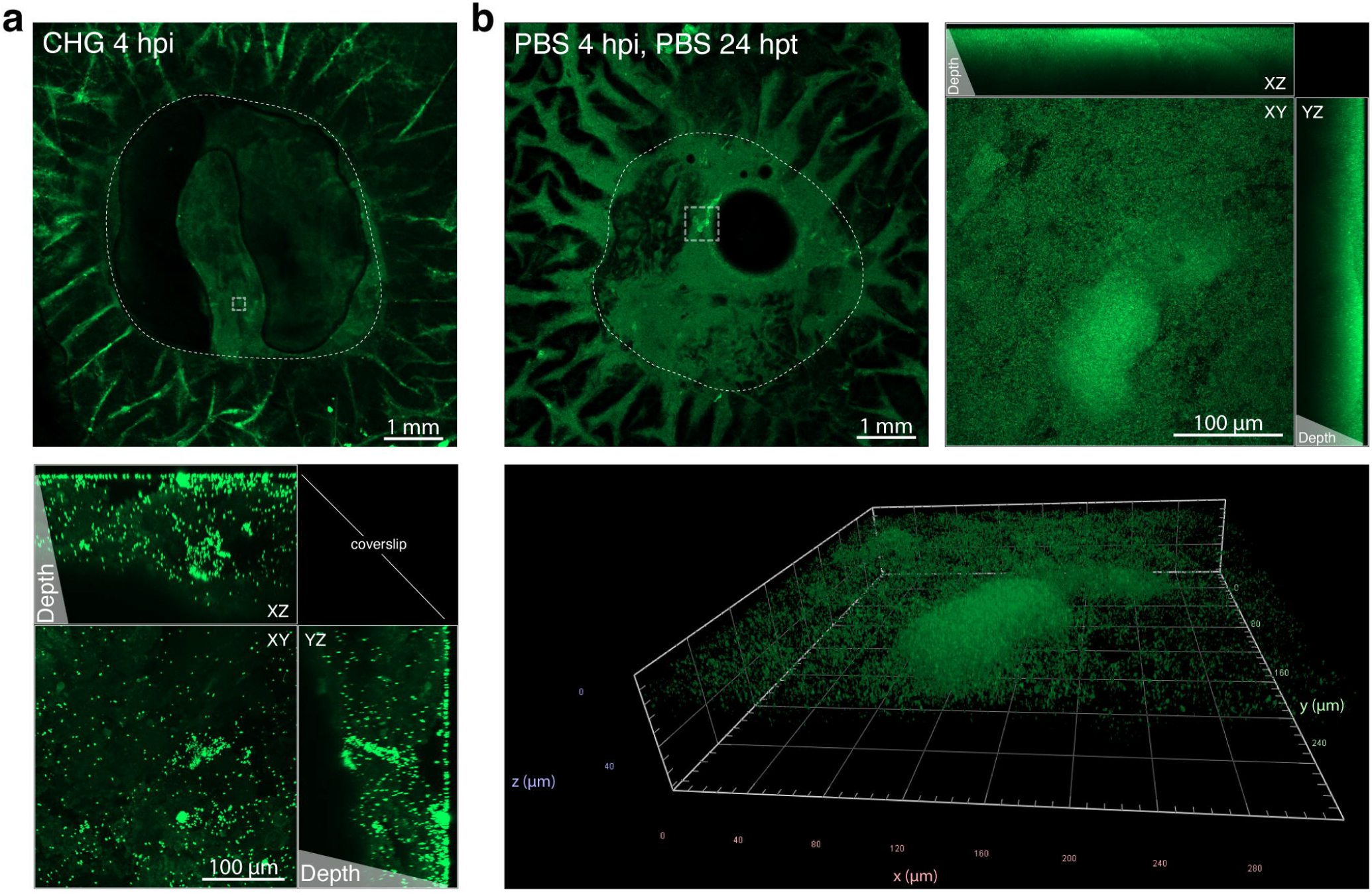
Confocal laser scanning microscopy shows bacterial aggregates and single-cells deep within wound bed. Live imaging of *ex vivo* wounds at 24 h after treatment. Circular outlines indicate the wound edge. Square outlines represent region magnified. Dark areas within the wound bed are imaging artifacts from air bubbles. (a) PBS treatment at 4 hpi, PBS treatment at 24 hpt. Maximum intensity projection shows a large aggregate of GFP-expressing *Pseudomonas aeruginosa*. (b) CHG treatment at 4 hpi. Maximum intensity projection shows single cells and small aggregates of bacteria found dispersed deep within the tissue.

## DISCUSSION

Choosing an appropriate wound cleansing agent requires balancing the desired antimicrobial efficacy with cytotoxic effects (Ratliff, 2014; Smith, 2005). We used an *ex vivo* human skin excisional wound model to investigate the localization of antiseptic-induced loss of tissue viability, and found that chlorhexidine gluconate (CHG) cytotoxicity persists and spreads over time. In parallel, we developed a wound contamination model, showing that inoculation of wound tissue with as little as ∼100 cells is sufficient to allow robust growth and accumulation of microbial biomass by the pathogen *Pseudomonas aeruginosa*. CHG imparts antiseptic activity immediately post-treatment, with effects lasting up to 24 hpt. However, we observe a loss of antimicrobial activity when applied daily, as is often done clinically. This suggests that antiseptic efficacy is short-lived and overcome by microbial growth dynamics within the wound environment.

We established a wound contamination model using ∼100 cells of *P. aeruginosa* per wound to evaluate antiseptic efficacy and microbial biofilm architecture with temporal resolution (Fig. 1). This is in contrast to wound infection models that inoculate at 10^5^ CFUs or higher (Johani et al., 2018; Wolcott et al., 2010; Yoon et al., 2019). Further, classical *in vitro* models often expose established microbial biofilms in environmental conditions that are divorced from the host context. For example, host-derived biofilm components such as fibrin and leukocyte-associated proteins can alter biofilm composition and structure (Besser et al., 2020; Kwiecinski et al., 2016; Nett et al., 2015; Rembe et al., 2020), with potential effects on antiseptic susceptibility, virulence and pathogenesis. Our model provides a clinically-relevant environmental context for microbial growth.

We evaluated the cytotoxicity of five clinically-used wound cleaning agents (saline, povidone iodine [PVI], Dove® soap, Dial® soap, and CHG) on an *ex vivo* model of human skin excisional wounds. Classical models to evaluate cytotoxicity are typically performed *in vitro* and lack structural components and cellular polarization of full-thickness skin (Borenfreund and Puerner, 1985; McLaughlin et al., 2020; Muller and Kramer, 2008; Teepe et al., 1993). We found that Dial® soap and CHG significantly reduced metabolic activity of the biopsies, while all treatments except saline resulted in localized cytotoxicity on the wound bed (Fig. 2b, c). In addition, the impact of CHG on the epidermal cells at the wound edge appears irreversible and cell death progresses to the epidermis lateral to the treated wound (Fig. 3). We hypothesize that antiseptic cytotoxicity is affected by contact and diffusion gradients of the antiseptic that occur throughout the heterogenous structure of a wound, which may also impact antiseptic efficacy on contaminating microbes. Given the ubiquitous use of CHG in clinical practice, these findings warrant further study to determine if a change in practice of CHG in wound care and in preoperative surgical preparations is necessary to prevent deleterious effects on wound healing.

We observe that CHG treatment is efficacious early on during the infection process as we were unable to detect bacterial counts immediately post-treatment (Fig. 4b). However, the bacterial population rises to an average of ∼10^5^ CFU/bisect by 24 hpt (Fig. 2a). Although the quantity of *P. aeruginosa* counts at 24 hpt remain significantly lower than the controls, these data suggest that within our model, colonization with less than 50 CFUs (the limit of detection) is permissive to allow infection to proceed. By the time bacterial bioburden reaches ∼10^5^ CFU, a second application of CHG, mimicking a clinical treatment schedule, does not have measurable effects on microbial growth. All treatments at 24 hpt resulted in bioburdens of > 10^8^ CFU/bisect (Fig. 4c). This is likely due to reduced antiseptic efficacy against microbial biofilms that may be forming in the wound environment, in contrast to activity against *in vitro* biofilms or planktonic cells in the early stages of wound contamination and colonization (Johani et al., 2018; Karpiński and Szkaradkiewicz, 2015; Vestby and Nesse, 2015). Notably, PVI treatment did not lower wound bioburden, despite robust *in vitro* efficacy (Barreto et al., 2020; Capriotti et al., 2018; Hoekstra et al., 2017; Johani et al., 2018)

Using SEM, we showed extensive surface colonization of the wound bed corresponding to high bacterial bioburden (Fig. 5). We observed a dense biofilm comprised of bacterial cells and extracellular matrix, which matured into mushroom-like aggregates (Fig. 6a). The spatial and physical structure of microbial biofilms are important for their virulence and pathogenesis (Nadell et al., 2016). Interestingly, wounds treated with CHG at 4 hpi showed no visible bacterial cells on the surface of the wound bed using SEM (Fig. 5c). However, CLSM revealed single bacterial cells and small aggregates dispersed within the tissue (Fig. 6b). Our model allows for evaluating spatial heterogeneity of microbes within the wound. For example, *P. aeruginosa* has been reported to reside deep within patient wound tissue as compared to other wound pathogens such as *Staphylococcus aureus* (Fazli et al., 2009).

In conclusion, we present a clinically-relevant model for evaluating antiseptic cytotoxicity and efficacy, with the ability to resolve spatial localization and temporal dynamics of tissue viability and microbial growth. We anticipate that this model will bolster basic, translational, and pre-clinical studies in wound care by providing further insights into the complex interplay between host responses and microbial growth dynamics in the context of advanced wound care and increasing use application of topical antiseptics to control microbial bioburden.

## MATERIALS AND METHODS

### *Ex vivo* Excisional Wound Model

Human skin was obtained from patients undergoing elective reconstructive surgeries. The de-identified samples were exempt from the regulation of University of Wisconsin-Madison Human Subjects Committee Institutional Review Boards. The tissue was rinsed with PBS and partial-thickness wounds were made by puncturing the epidermis with a 6 mm biopsy punch and removing the entire epidermis and a portion of the dermis. A 12 mm biopsy punch was then used to make full-thickness biopsies with the wound. In the antiseptic antimicrobial efficacy, biopsies were placed into 12-well plates containing 3 mL of a DMEM-agarose gel (0.15:0.85 ratio of 1% agarose in PBS and Dulbecco’s modified Eagle medium [DMEM] supplemented with 10% fetal bovine serum [FBS]). In the antiseptic cell viability studies, biopsies were placed onto a fine mesh insert in p100 plates to raise the tissue to the air-liquid interface in media containing 10 mL of DMEM (Gibco, Thermo Fisher, Waltham, MA) supplemented with 10% FBS (Gibco, Thermo Fisher, Waltham, MA), 0.625 µg/mL Amphotericin B (Thermo Fisher, Waltham, MA), and 100 µg/mL of Pen/Strep (Thermo Fisher, Waltham, MA). Biopsies were incubated at 37°C with 5% CO_2_ overnight before inoculation or treatment, and were transferred to a new medium every 48 h.

### *Ex vivo* Wound Colonization Model

Inoculums were prepared by suspending colonies of *Pseudomonas aeruginosa* PA1 (GFP plasmid with carbenicillin resistance) grown overnight at 37°C on tryptone soy agar (TSA) plates supplemented with 200 μg/mL carbenicillin in PBS and diluted to a cell density of 1 × 10^4^ CFU/mL. Wounds were inoculated within 24 to 48 h of tissue collection with 10 μL of inoculum for a final cell density of 1 × 10^2^ CFU/bisect. Following 4 h of incubation, wounds were treated with antiseptics (see below), rinsed, incubated for 24 h, and then processed for microscopy (see below) or bisected and processed for viable cell enumeration. A subset of PBS-and CHG-treated biopsies was immediately processed after treatment at 4 hpi, or treated for a second time at 24 hpt and incubated for 24 h before processing. All bisects were vortexed in 1 mL PBS with 5% Tween 80 and 0.6% sodium oleate (as CHG neutralizer) with 0.2 g of 1 mm sterile glass beads for 10 min at full-speed on a Vortex-Genie 2 (Scientific Industries, Bohemia, NY) before serial dilution and spot plating 20 μL on TSA plates with no antibiotic supplementation.

### Antiseptic Treatment

*Ex vivo* wound tissue biopsies were treated with 5 antiseptics including PBS, PVI, Dial® and Dove® soaps, and CHG. Dial® and Dove® soaps were diluted 1:1 in sterile water 24 h before treatment and allowed to mix. Wound cleansing solutions were applied onto the wound by gently blotting with sterile cotton swabs around the wound edge until the solution pooled on the wound bed. The treatment was left on for 30 min before gently rinsing twice with PBS. For cell viability studies, the biopsies were incubated for an additional 24 h before the MTT assay and LDH staining were performed. To determine whether the cytotoxicity induced by antiseptics can recover over time, a subset of PBS-and CHG-treated biopsies were cultured at 37°C and 5% CO_2_ for up to 14 days. Tissues were harvested on day 1, 3, 7 and 14 and processed for LDH staining (see below).

### Tissue Metabolic Activity Assay

At 24 hours post-treatment, cell viability of treated tissues was quantified using a tetrazolium-based (MTT) assay. Briefly, each bisect was rinsed in PBS and placed in an individual well of a 6-well plate with 2 mL MTT solution (2 mg/mL, Invitrogen) in each well. The 6-well plates were placed on a rotating plate and incubated at 37°C at 100 rpm for 2 h. After aspiration of the remaining MTT solution, 4 mL DMSO was added to each well and incubated at 100 rpm at 37°C for 80 min. 200 µL aliquots of solution in each well were transferred to a 96-well plate with DMSO blank controls. The optical density of the solution was measured using a plate reader (FlexStation 3, Molecular Devices) at a wavelength of 540 nm. For the MTT assay, untreated tissue biopsies were immersed in boiling water for 30 min as a negative control, while the PBS-treated biopsies served as the positive control.

### Histological tissue processing

Tissue bisects were snap-frozen in Tissue-Tek optimum cutting temperature (OCT) compound (Sakura Finetek USA Inc, Torrance, CA) for cryo-sectioning into 8 µm sections before staining for lactate dehydrogenase activity (Gibson et al., 2017). Histological slide sections were examined under a Nikon Ti-S inverted microscope and scanned at 4x using a slide scanner (PathScan Enabler 5, Meyer Instruments, Houston, TX).

### Confocal Microscopy

Biopsies were mounted in glass-bottomed 60 mm petri dishes (14 mm opening; MatTek, Ashland, MA) and imaged on a Zeiss 780 confocal laser scanning microscope on the FITC channel using 5x and 40x objectives. Zeiss Zen software was used to analyze z-stacks and generate maximum intensity projections.

### Scanning Electron Microscopy

The following protocol was adapted from Horton et al. (2020). Briefly, *ex vivo* human skin wounds were rinsed with PBS and fixed overnight in 5 mL of 1.5% glutaraldehyde in 0.1 M sodium phosphate buffer (pH 7.2) at 4°C. Samples were rinsed, treated with 1% osmium tetroxide for 1 h, and then washed again. Samples were dehydrated through a series of ethanol washes (30%-100%) followed by critical point drying (14 exchanges on low speed) and were subsequently mounted on aluminum stubs with a carbon adhesive tab and carbon paint. Silver paint was applied around the perimeter for improved conductivity. Samples were left to dry in a desiccator overnight. Following sputter coating with platinum to a thickness of 20 nm, samples were imaged in a scanning electron microscope (Zeiss LEO 1530-VP) at 3 kV.

### Data and Statistical Analysis

Information regarding sample size and replication are described in the figure legends. All statistical analysis was performed using R (R Core Team, 2020). Multiple comparisons and estimation of mean differences between inoculation conditions for each microbe were evaluated using a one-way between subjects ANOVA with Tukey’s Honest Significant Differences test. We used an α level of 0.05 for all statistical tests.

## Abbreviations

CFU: colony forming unit
CHG: chlorhexidine gluconate
CLSM: confocal laser scanning microscopy
DMEM: Dulbecco’s modified Eagle’s medium
FBS: fetal bovine serum
GFP: green fluorescent protein
hpi: hours post-inoculation
hpt: hours post-treatment
LDH: lactate dehydrogenase
PBS: phosphate-buffered saline
PVI: povidone iodine
SEM: scanning electron microscopy

## DATA AVAILABILITY STATEMENT

Datasets related to this article are provided in spreadsheets in the Supplementary Data File.

## ACKNOWLEDGEMENTS

The authors would like to thank Nathan Holly, Edgar Ocotl, and Lily Meronek for technical assistance. The authors acknowledge use of facilities and instrumentation at the UW-Madison Wisconsin Centers for Nanoscale Technology (wcnt.wisc.edu) partially supported by the NSF through the University of Wisconsin Materials Research Science and Engineering Center (DMR-1720415). Figures include the icons Skin by Hermine Blanquart, Energy by Alice Design, Bacteria by Maxim Kulikov, Cotton Swab by Kid Kitaro, Petri Dish by BOYS, Pipette by Joseph L Elsbernd, and Skin by Fauzan Akbar, all from the Noun Project

This work was supported by grants from the Wisconsin Partnership Program and Shapiro Summer Research Program from the School of Medicine and Public Health, University of Wisconsin–Madison (A.L.F.G).

## AUTHOR CONTRIBUTIONS

